# Zones of Cellular Damage Around Pulsed-Laser Wounds

**DOI:** 10.1101/2021.05.28.446116

**Authors:** James O’Connor, Fabiha Bushra Akbar, M. Shane Hutson, Andrea Page-McCaw

## Abstract

After a tissue is wounded, cells surrounding the wound adopt distinct wound-healing behaviors to repair the tissue. Considerable effort has been spent on understanding the signaling pathways that regulate immune and tissue-resident cells as they respond to wounds, but these signals must ultimately originate from the physical damage inflicted by the wound. Tissue wounds comprise several types of cellular damage, and recent work indicates that different types of cellular damage initiate different types of signaling. Hence to understand wound signaling, it is important to identify and localize the types of wound-induced cellular damage. Laser ablation is widely used by researchers to create reproducible, aseptic wounds in a tissue that can be live-imaged. Because laser wounding involves a combination of photochemical, photothermal and photomechanical mechanisms, each with distinct spatial dependencies, cells around a pulsed-laser wound will experience a gradient of damage. Here we exploit this gradient to create a map of wound-induced cellular damage. Using genetically-encoded fluorescent proteins, we monitor damaged cellular and sub-cellular components of epithelial cells in living *Drosophila* pupae in the seconds to minutes following wounding. We hypothesized that the regions of damage would be predictably arrayed around wounds of varying sizes, and subsequent analysis found that all damage radii are linearly related over a 3-fold range of wound size. Thus, around laser wounds, the distinct regions of damage can be estimated after measuring any one. This report identifies several different types of cellular damage within a wounded epithelial tissue in a living animal. By quantitatively mapping the size and placement of these different types of damage, we set the foundation for tracing wound-induced signaling back to the damage that initiates it.

## Introduction

The field of wound repair and regeneration has long sought to connect the signals emanating from wounds to the behavioral changes undertaken by cells around the wound. In response to wounds, tissue-resident cells transition from a stationary and non-proliferative state to a migratory or proliferative state, (Antunes et al., 2013; Benink and Bement, 2005; Bosch et al., 2005; Capilla et al., 2017; Carvalho et al., 2014; Danjo and Gipson, 2002; Hunter et al., 2015; Martin and Lewis, 1992; Matsubayashi and Millard, 2016; Park et al., 2017; Radice, 1980; Rämet et al., 2002; Stramer et al., 2005; Wood et al., 2002) whereas immune cells migrate from outside the tissue to clear debris and fight infection by pathogens entering through the wound (Fauvarque and Williams, 2011; Luster et al., 2005; Razzell et al., 2013; Thuma et al., 2018). Ultimately, the signals emanating from wounds must be derived from the damage itself; however, only a few studies have characterized the damaged tissue on a cellular/sub-cellular level to understand the distinct types of damage created by wounds (Hellman et al., 2008; Hutson and Ma, 2007; McNeil and Steinhardt, 1997; Welch et al., 1991). Importantly, our previous studies have found that multiple signaling pathways are initiated around the same time within the same wound, and further, that distinct types of cellular damage initiate each signaling pathway. Specifically, cell lysis leads to the release of cellular proteases, which cleave and activate cytokines in the vicinity of the wound, which in turn signal to surrounding cells through G-protein coupled receptors (O’Connor et al., 2020). However, even before cytokine signaling is evident, cells with a different type of damage – torn plasma membranes – initiate calcium signaling, as extracellular calcium floods in through damaged membranes and out through gap junctions to undamaged neighboring cells (Shannon et al., 2017). Together, these studies suggest that understanding the origin of wound-induced signaling requires identifying and categorizing different types of cellular damage induced by wounding. In this study, we attempt to solve this problem by classifying various zones of damage around epithelial wounds visualized using genetically-encoded fluorophores, and mathematically defining the relationships between these zones of damage to create a map of the characteristic cellular damage experienced by a tissue following a wound.

It may be impossible to develop a useful map of cellular damage around ordinary trauma wounds experienced in the natural world, as the types of damage would be irregularly arrayed and would not be sufficiently reproducible to identify patterns. However, researchers often employ laser-induced wounding, which has advantages compared to other methods of wounding because of its ability to make reproducible, radially-symmetric, aseptic wounds in a tissue while simultaneously imaging (Antunes et al., 2013; Benink and Bement, 2005; Davenport et al., 2016; Enyedi and Niethammer, 2015; Gault et al., 2014; Hunter et al., 2018; Razzell et al., 2013; Shannon et al., 2017; Stramer et al., 2005; Wood et al., 2002). Two types of lasers are used to damage tissues–continuous wave lasers and pulsed-lasers–which make very different types of cellular damage. A continuous wave laser delivers energy to the sample on the order of milliseconds to seconds, and in doing so delivers a high amount of thermal energy to both the ablated region and the surrounding tissue, causing burning and/or explosive ejection of cellular debris from the sample (Welch et al., 1991). In contrast, a pulsed-laser delivers pulses of energy on the order of picoseconds to microseconds, dramatically decreasing the thermal energy transfer to the surrounding tissue (Troutman et al., 1986; Welch et al., 1991). A pulsed-laser superheats only the ablated region, vaporizing it into a bubble of gas known as a cavitation bubble, which expands and contracts within microseconds, causing mechanical (but not thermal) damage to the surrounding tissue (Hutson and Ma, 2007; Leeuwen et al., 1995; Venugopalan et al., 2002; Welch et al., 1991). Although continuous lasers were often used through the 1980s, pulsed-lasers have become more widely used over the last three decades because of the shorter thermal event, delivering energy so rapidly as to destroy and remove hot tissue before the heat can be transferred, thus reducing thermal damage to the surrounding tissue and removing burn damage as a confounding variable in studying wound signaling and repair (Leeuwen et al., 1995; Troutman et al., 1986). Although other methods of wounding have been utilized in wound repair studies, namely puncture (Burra et al., 2013; Galko and Krasnow, 2004; Xu and Chisholm, 2011), pinch (Ramos-Lewis et al., 2018; Wu et al., 2009), scratch wounding (Shabir and Southgate, 2008), or complete amputation (Martin et al., 1994; Yoo et al., 2012), these methods can introduce experimentally-induced variability that hinders reproducibility.

Previously, our labs used pulsed-laser wounding on an epithelial monolayer in living *Drosophila* pupae to understand wound-induced repair signals, specifically how calcium is increased in the cytosol of cells around wounds (Shannon et al., 2017). Unexpectedly we found that the laser-induced cavitation bubble damages the plasma membranes of epithelial cells around the wound margin, creating microtears that allow extracellular calcium to flood into damaged cells within milliseconds of wounding. To study this phenomenon, we varied the laser energy and found that cavitation bubble area matched the areas of extracellular calcium entry, membrane depolarization, and extracellular dye entry, three indicators that plasma membrane integrity was lost in the tissue damaged by the cavitation bubble (Shannon et al., 2017). Many of the cells with plasma membrane damage were able to repolarize and survive, demonstrating that not all damaged cells die. Other reports have demonstrated loss of plasma membrane integrity of cells around wounds made by multiple types of injury (Davenport et al., 2016; McNeil and Ito, 1989, 1990; McNeil and Steinhardt, 1997; Shabir and Southgate, 2008). Indeed, in these studies many of these cells with plasma membrane damage survived and were able to restore membrane integrity (Davenport et al., 2016; McNeil and Steinhardt, 1997). Thus, these previous studies show there are *at least* two populations of cells that experience damage following a wound: cells near the center of the wound that are damaged so severely that they are destroyed and cells distal from the center of the wound that are damaged but ultimately survive and participate in the repair process. In this study, we identify several more types of cells that are damaged by laser wounds; specifically, we monitored initial laser rupture, cell lysis, nuclear membrane damage, plasma membrane damage, chromatin disruption, Ecadherin loss, and a calcium expansion outward from the wound site. The severity of damage was inversely correlated with radius from the wound center, and by varying wound size, we find that these regions of damage are arrayed in predictable and reproducible patterns around the wound.

## Results

### Live imaging wounded Drosophila pupae

We chose *Drosophila melanogaster* (fruit flies) for wounding studies because they are genetically tractable, with numerous existing strains expressing genetically-encoded fluorescently-tagged proteins easily visualized during live imaging. We analyzed wounds in the pupal notum, the epithelial tissue on the dorsal thorax of the pupa, easily accessible after partially removing the pupal case (Antunes et al., 2013) (Fig. 1A). Wounds were administered by pulsed-laser ablation, allowing simultaneous wounding and imaging *in vivo*. Because the pupa is stationary, it can be easily mounted, wounded, and imaged for extended periods of time (hours to days) and still survive to adulthood.

**Figure 1:**
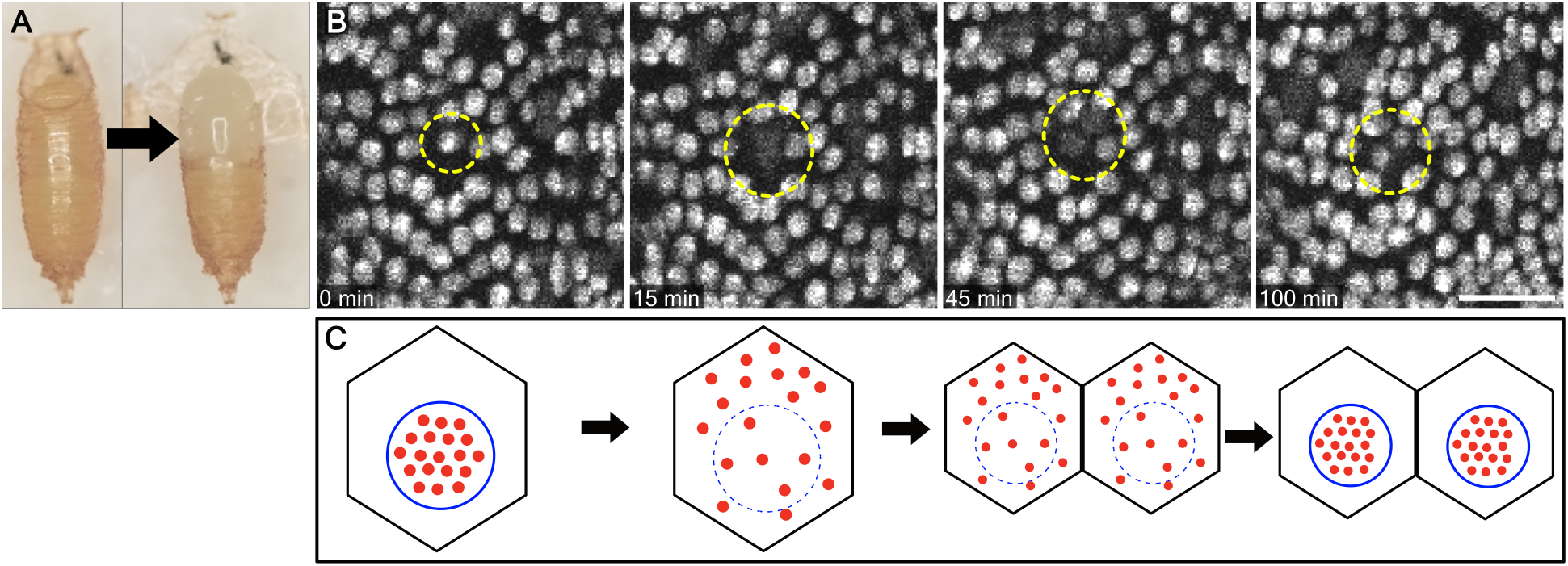
mCherry.NLS reports nuclear envelope integrity. (A) An intact *Drosophila* pupa (left) and one prepared for imaging with the pupal case partially removed (right), exposing the pupal notum. (B) Interphase nuclei in unwounded tissue are labeled by mCherry.NLS driven by the *Gal4/UAS* system. During mitosis, the fluorescence signal evident at 0 min disperses when the nuclear envelope breaks down, seen within circles in panels at 15 min and 45 min, before reappearing as two distinct/punctate nuclei at 100 min. (C) A schematic showing the diffusion of mCherry.NLS out of the nucleus during mitosis and its concentration in nuclei again after the nuclear envelope reforms in the daughter cells. Thus, the localization of mCherry.NLS reports nuclear membrane integrity. Scale bar = 25 µm.

Using the *Gal4/UAS* system (Brand and Perrimon, 1993), we drove expression of the fluorescent protein mCherry with a nuclear localization signal (mCherry.NLS) in the pupal notum to visualize live epithelial nuclei. In unwounded animals, we observed that the punctate mCherry.NLS frequently became diffuse during normal cell division, then re-appeared as two separate punctae (Fig. 1B). mCherry.NLS is a freely-diffusible fluorescent protein carrying a nuclear localization signal, and this pattern indicates that mCherry.NLS diffused into the cytosol during nuclear membrane breakdown (Fig. 1C).

Upon pulsed-laser ablation, the mCherry.NLS was lost at the wound center within the first two frames after ablation (usually ∼2 seconds). Because the fluorophore diffused away from both the nucleus and the plasma membrane to leave the tissue (orange circle, Fig. 2A’), this loss of fluorescence indicates cell destruction and rupture. We term this immediate loss of mCherry the **region of laser rupture**. This region included the area nearest the laser’s focal point, which was immediately vaporized, and it is much smaller than the area of the cavitation bubble (see below). In the wounds made for this study, the size of laser rupture ranged between 10 and 30 µm in radius.

**Figure 2:**
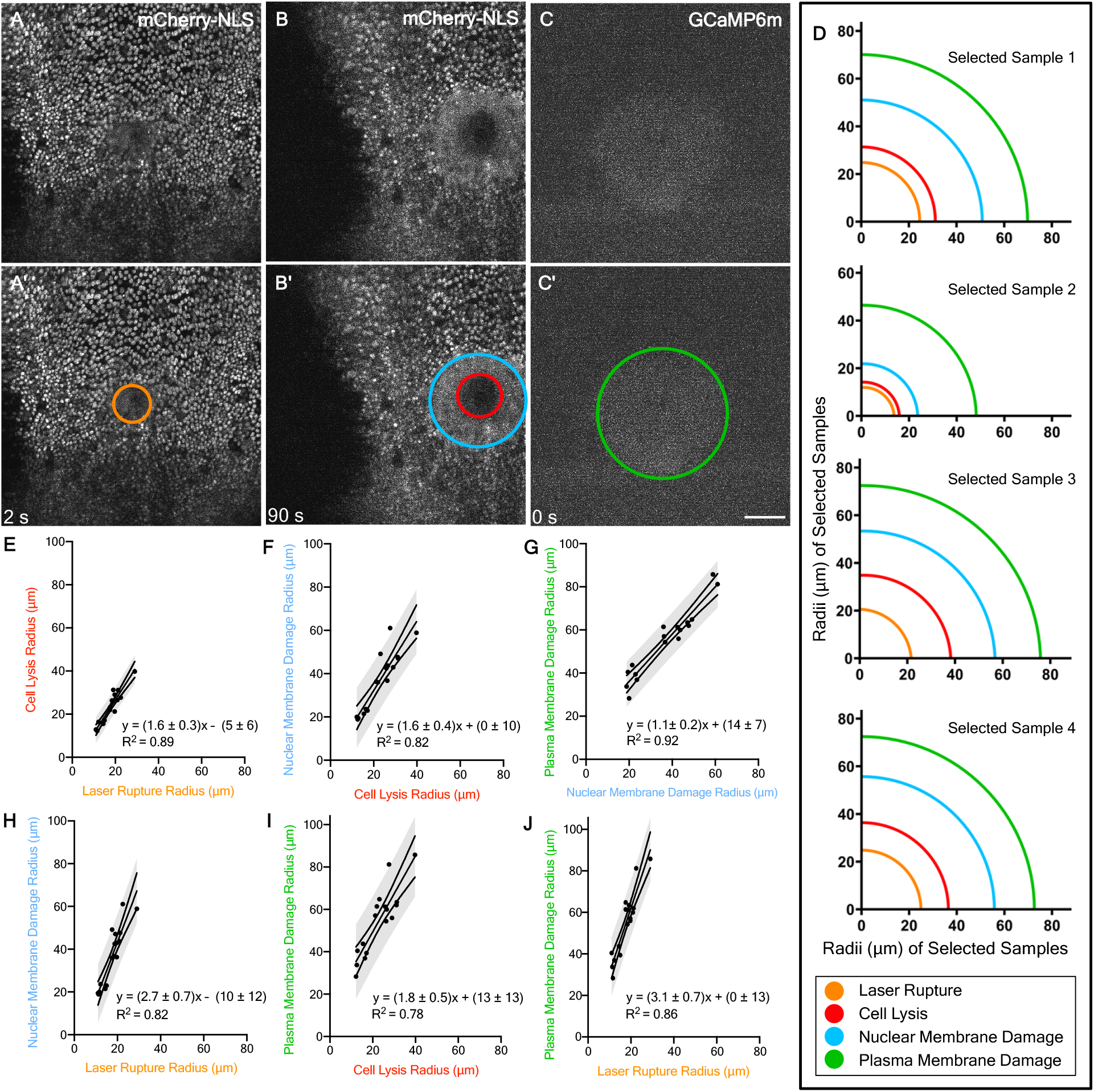
Several zones of damage are evident around pulsed-laser wounds. (A) The region of laser rupture is observed as the area of disrupted mCherry.NLS within the first two frames after wounding (here 2 seconds), annotated with an orange circle in A’. (B) The region of cell lysis is observed as the complete loss of mCherry.NLS at 90 seconds, (red circle in B’) and the region of nuclear membrane damage is observed as a diffuse, non-nuclear mCherry.NLS signal (light blue circle in B’), maintained within the cells by the plasma membrane at 90 seconds after wounding. (C) The region of plasma membrane damage is observed as an increase in cytoplasmic calcium levels immediately after wounding (green circle in C’, as reported by GCaMP6m, a fluorescent calcium indicator, in the first frame after wounding (here 0 seconds). In this region, the wound creates microtears in the plasma membrane, allowing immediate influx of extracellular calcium. (D) Actual measurements of radii in four different samples demonstrates a consistent relationship between these regions; as any one of these regions increases in radius, the others likewise increase in radius. Panels E-J quantify this trend with 95% prediction bands. Scale bar = 50 µm.

Within 90 seconds after pulsed-laser ablation, the mCherry.NLS signal changed in two distinct ways. First, the region devoid of mCherry signal increased in size, indicating that even after ablation, cells continue to lyse (compare Fig. 2A,B). No fluorescence signal returned to this region until late in the repair process when distal cells migrated in to repair the wound. This larger area of cell death we term **region of cell lysis** based on the complete loss of mCherry.NLS (red circle, Fig. 2B’). Both the initial laser energy and the ensuing cavitation bubble may contribute to cell lysis in this area.

Second, beyond the cell lysis area, the mCherry.NLS became diffuse, no longer confined to the nucleus but still remaining in the tissue (light blue circle, Fig. 2B’), similar to its appearance during mitosis (Fig. 1B). Interestingly, later in the repair process the mCherry signal returned to the nucleus in some of these cells, indicating that some of these cells survive. We concluded that these cells have undergone damage to the nuclear membrane, causing the previously nuclear-localized mCherry to be released from the nucleus into the cytosol (Fig. 2B). We term this the **region of nuclear membrane damage**.

To understand how these zones of damage compared with other types of damage we had already characterized, we compared the mCherry.NLS patterns with those of GCaMP, a ubiquitously expressed genetically encoded cytoplasmic calcium reporter. Our previous work demonstrated that within the first frame after pulsed-laser ablation, cells near the wound exhibit an increase in GCaMP fluorescence. This region of immediate GCaMP fluorescence indicates the region where the plasma membrane is torn by the laser-induced cavitation bubble, allowing the rapid influx of extracellular calcium (Shannon et al., 2017). The cavitation bubble expands and collapses within microseconds, and extracellular calcium enters through microtears in the plasma membrane within ∼10 milliseconds, increasing its concentration over the next many seconds as more calcium enters the cell (Shannon et al). Thus, the immediate GcaMP signal indicates the **region of plasma membrane damage** (green circle, Fig. 2C’). Although the GCaMP signal is still dim in the first frame after wounding, the first frame provides the most accurate measure of the area of plasma membrane damage because the calcium diffuses outward to neighboring cells through gap junctions over the next 15-20 seconds, enlarging the area of GCaMP6m fluorescence beyond the region of plasma membrane damage (Shannon et al., 2017).

After measuring the radii of these four well-defined regions in multiple samples with varying wound sizes, we found the four regions maintained the same ascending size order within every sample: Laser rupture < cell lysis < nuclear membrane damage < plasma membrane damage (Fig. 2D). To determine whether there was a consistent relationship between the radii of each region, we analyzed how each of these four regions related to one another in wounds of various sizes and graphed the six relationships (Fig. 2E-H). This analysis revealed that the radius of each zone was linearly correlated with the radius of each other zone, with an R^2^ value for each of the trendlines lying between 0.78 and 0.92. Importantly, these results show that each laser-induced wound creates reproducible regions of damage around a single wound, and all of these regions can be reasonably estimated after measuring the radius of a single one. Not only were adjacent zones of damage related linearly, but also the linear relationship was maintained even between laser rupture and plasma membrane damage, the smallest and the largest zones of damage respectively (Fig. 2J).

To better understand the nuclear membrane damage, we co-expressed mCherry.NLS and a GFP-tagged histone variant, His2Av-GFP. Unlike mCherry.NLS, the GFP-tagged histone is a fusion protein incorporated into chromatin, regardless of cell cycle stage or nuclear envelope integrity. Therefore, we could track nuclear destruction independently of nuclear membrane damage. At 5 minutes after wounding, there were many intact, well-ordered, His2Av-GFP-containing nuclei within the region of nuclear membrane damage, confirming that the nuclear membrane, but not necessarily chromatin, are damaged in this region (Fig. 3A-C). However, toward the center of the wound there were some misshapen punctae of His-GFP (Fig. 3A-C). We term this the **region of chromatin disruption**, and we measured it and compared it to the regions of cell lysis and nuclear membrane damage (Fig. 3A-C). We found that the chromatin disruption region was much smaller than the nuclear membrane damage region (Fig. 3D) and was similar in size to the region of cell lysis (Fig. 3E). These results demonstrate that even after cell lysis, the His-GFP and chromatin do not freely diffuse. Further, they raise the possibility that lysis and chromatin disruption are related. Though not the best-fit equation, we also checked the fit of y=x between the region of chromatin disruption and cell lysis and found the R^2^ was 0.84, which shows the two regions indeed correlate very well with each other.

**Figure 3:**
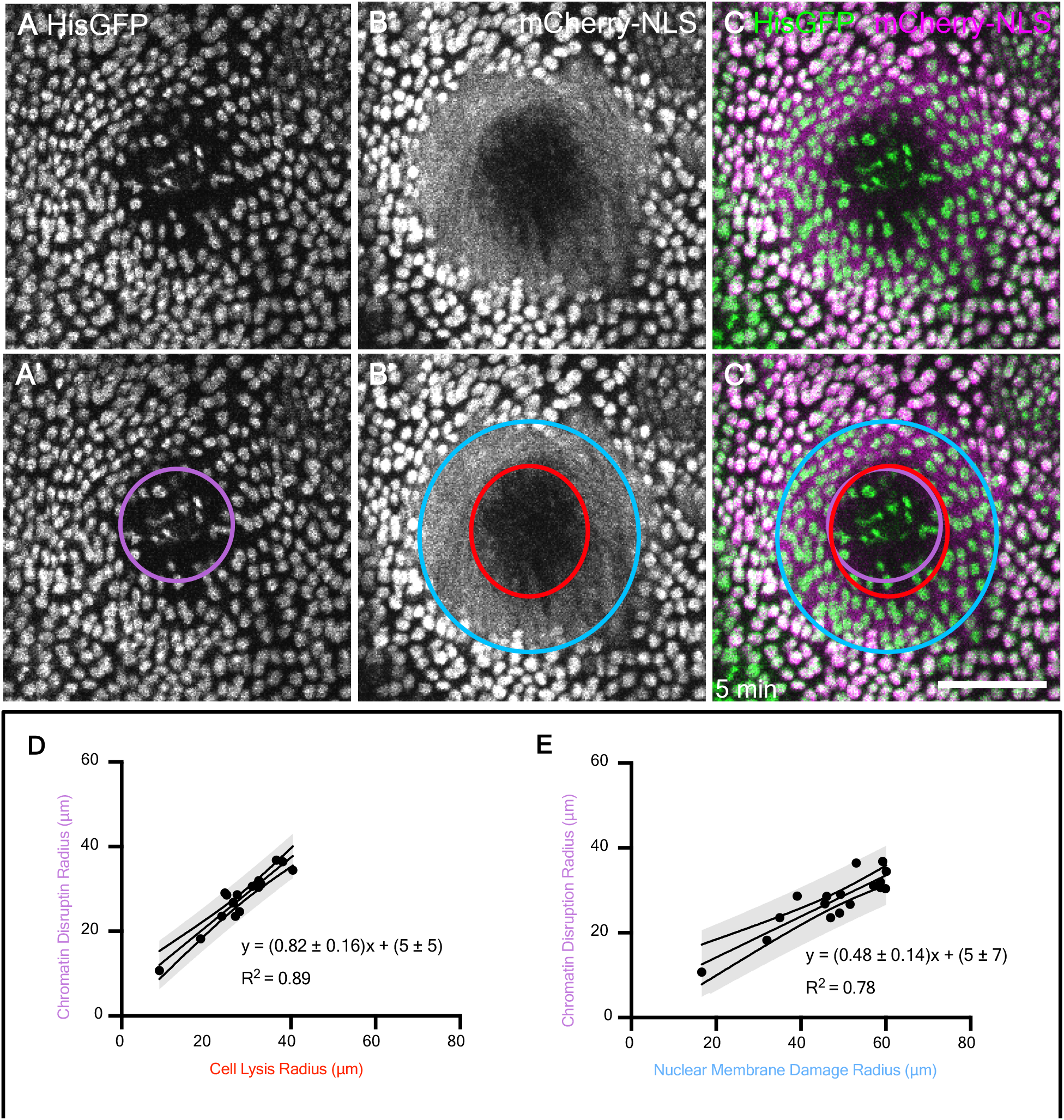
Histones reveal that within the region of nuclear membrane damage, some nuclei have intact chromatin whereas nuclei closer to the center have disrupted chromatin. (A) His2Av-GFP, which labels histones in chromatin, reveals a damage region with misshapen chromatin and decreased fluorescence, indicated with a purple circle in A’. (B) mCherry.NLS in the same wound reveals the region of nuclear membrane damage, (blue circle in B’) and the region of cell lysis (red circle in B’). (C) The overlay of His2Av-GFP with mCherry.NLS shows that the region of chromatin disruption is much smaller than the region of nuclear membrane damage, but similar in size to the region of cell lysis. (D-E) The relationships between the region of chromatin disruption and the regions of nuclear membrane damage (D) and of cell lysis (E) are linear. The 95% confidence interval is indicated. Scale bar = 50 µm.

To monitor individual cells in addition to individual nuclei, we co-expressed the mCherry.NLS fluorescent marker with GFP-tagged Ecadherin to visualize nuclei and cell borders simultaneously. Interestingly, a new region of damage was evident after wounding as a region of **immediate Ecadherin loss**. This region was evident starting at the first frame after wounding, although we often measured it at 5 min after wounding as it did not change during this time (Fig. S1). The Ecadherin labeling along cell borders appeared disrupted within a radius smaller than the region of nuclear membrane damage (yellow and light-blue circles respectively, Fig. 4A-C,E), and surprisingly, smaller than the region of cell lysis where cells had completely lost mCherry.NLS signal (yellow and red circles respectively, Fig. 4A-D). Ecadherin forms stable complexes with the adherens junctions that may be able to withstand damage such as cell lysis and release of cytoplasm.

**Figure 4:**
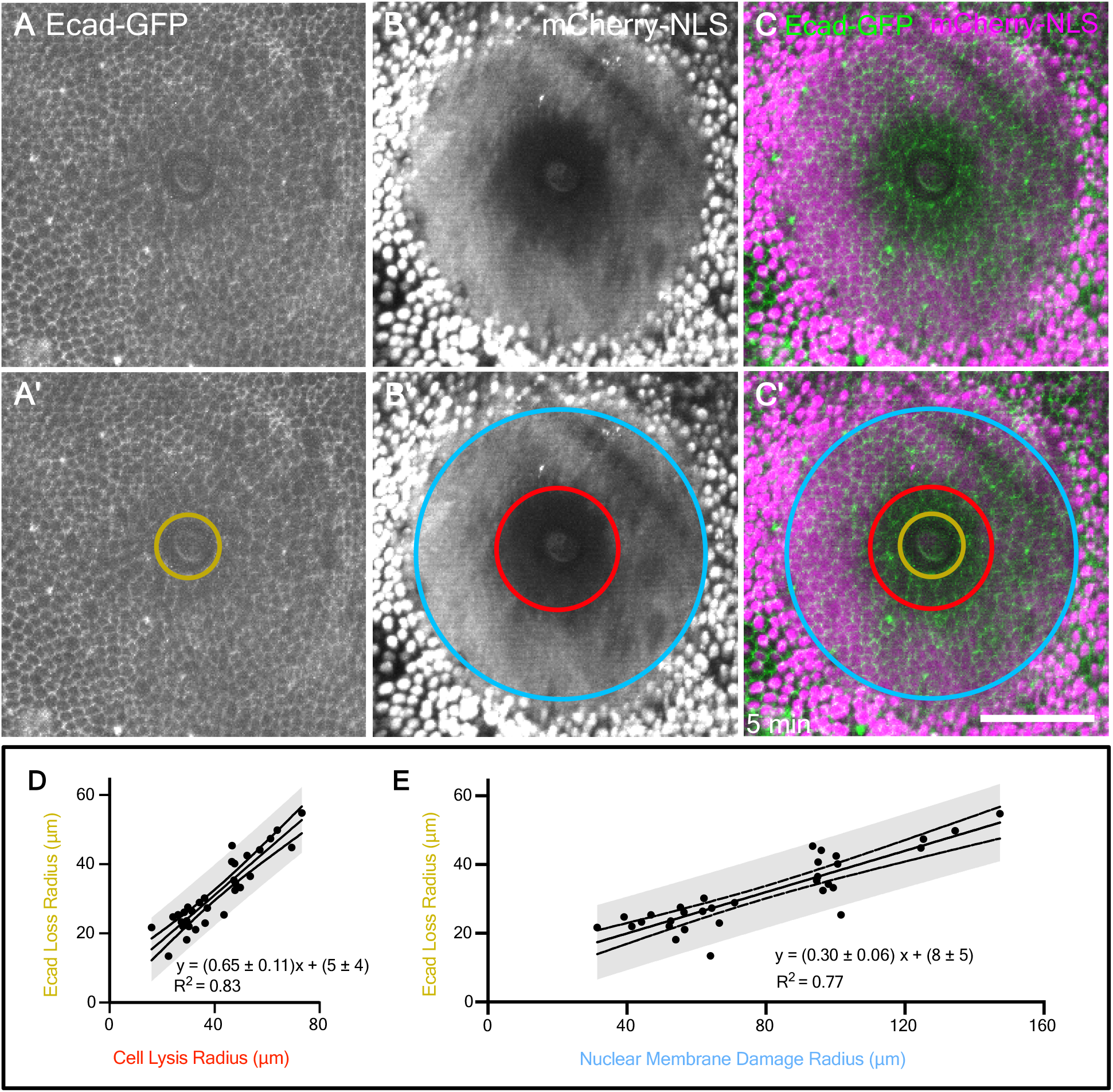
Cell borders are disrupted following wounding. (A) Genetically encoded Ecadherin-GFP marks cell borders and reveals damage around a pulsed-laser wound. The region of immediate Ecadherin-GFP loss is indicated with a gold circle in A’. (B,C) The region of immediate Ecadherin loss is distinct from and within the regions of cell lysis (red circle) and nuclear damage (blue circle). (D) The relationship of the radius of cell lysis to the radius of immediate Ecad loss is linear. (E) The relationship of the radius of nuclear damage to the radius of immediate Ecad-loss is linear. 95% prediction interval is indicated in D,E. Scale bar = 50 µm.

To determine the fate of cell borders around lysed cells, we imaged pupae co-expressing Ecadherin-GFP and mCherry.NLS for 90 minutes following wounding. Cell borders that initially appeared intact started breaking down around 30 minutes and were gone by 90 minutes (Fig. S1, Movie 1), disassembling throughout the region of cell lysis more or less simultaneously. The region where Ecad was dismantled (Fig. S1, yellow asterisks) spanned the entirety of the cell lysis region determined by mCherry.NLS (Fig. S1, red circle) as well as one to two cell diameters into the region of nuclear membrane damage (Fig. S1, light blue circle). This indicates that within the region of lysis, although the Ecad/adherens junctions appear stable five minutes after wounding, they represent the carcasses of dead/dying cells that do not survive the repair process.

The last damage-induced landmark we analyzed was the progression of the cytoplasmic calcium signal outward from the wound within the first seconds after wounding, measured as an increase in the radius of GCaMP fluorescence. After extracellular calcium enters cells through plasma membrane damage, that initial influx of calcium expands outward to neighboring cells through gap junctions within 15 seconds following wounding. We have termed this the **first calcium expansion**, so called because there is an independent second calcium response later (O’Connor et al., 2020). Although the first calcium expansion does not represent a type of cellular damage, it may represent damage-induced information transmitted to nearby cells, and we were interested to see how it compared in size to the damage itself. We compared the radius of the first calcium expansion to the radius of plasma membrane damage and found that it was positively correlated with a trendline of slope near 1, but with a y-intercept of 19 µm (Fig. 5). This linear relationship suggested that regardless of wound size, the initial influx of calcium travels about 20 µm outward to the next 2-4 neighboring cells, which experience this wound induced calcium signal even though they are ostensibly undamaged.

**Figure 5:**
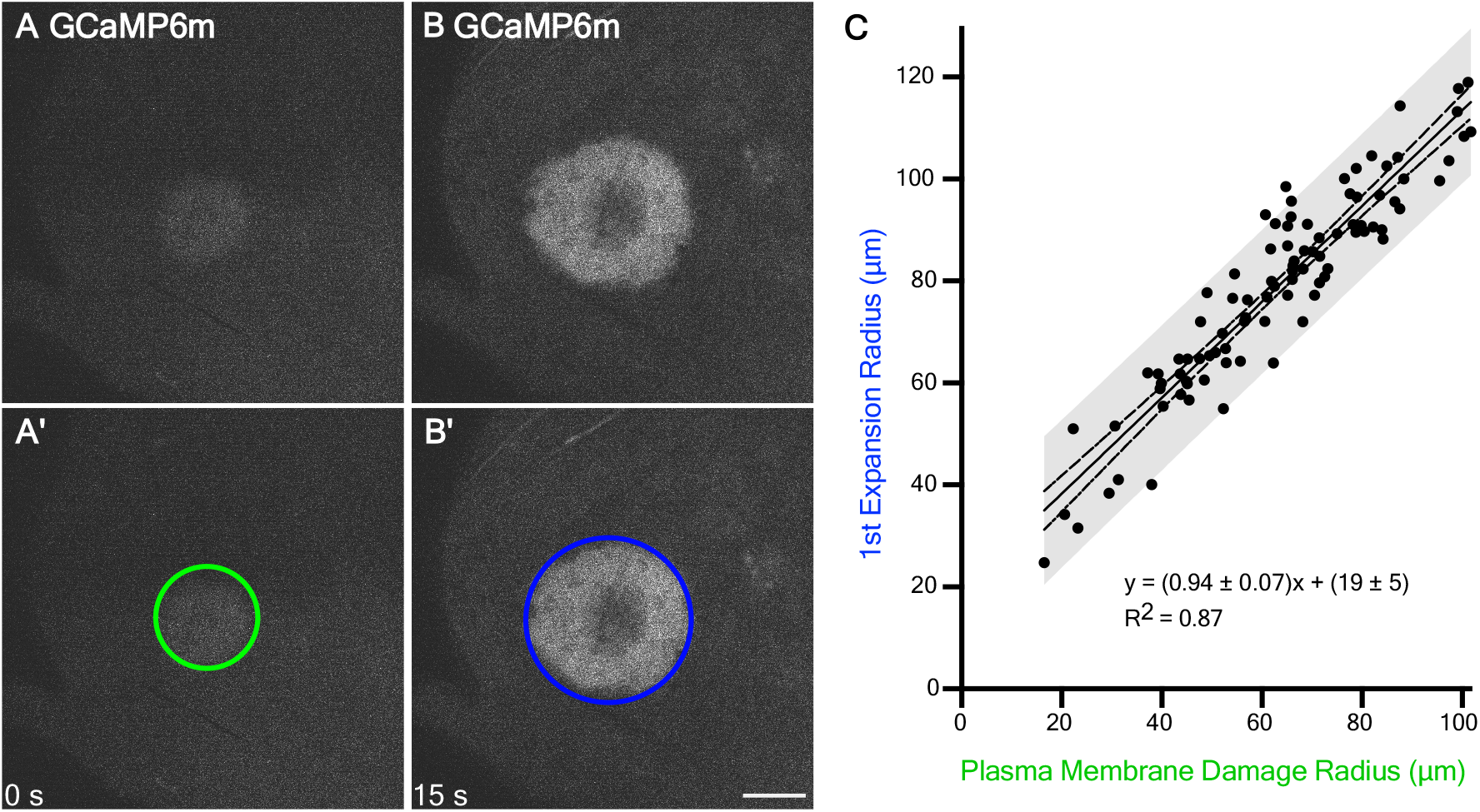
The first expansion of calcium extends about 20 µm from the region of plasma membrane damage, independent of wound size. (A) The influx of calcium immediately after wounding identifies cells with plasma membrane damage. This region is observed by an immediate increase in GCaMP fluorescence, indicated by a green circle in A’. (B) The calcium radius increases over the next ∼15 seconds (Shannon et al., 2017), and the maximum first expansion region is indicated with a dark blue circle in panel B’. The relationship between the plasma membrane radius and the first expansion radius is linear. Because the slope is ∼1, the y-intercept of 19 indicates that the 1^st^ expansion radius is expected to be ∼20 µm larger (∼2 to 4 cell diameters) than the radius of plasma membrane damage, regardless of wound size or laser energy. Scale bar = 50 µm.

**Figure 6:**
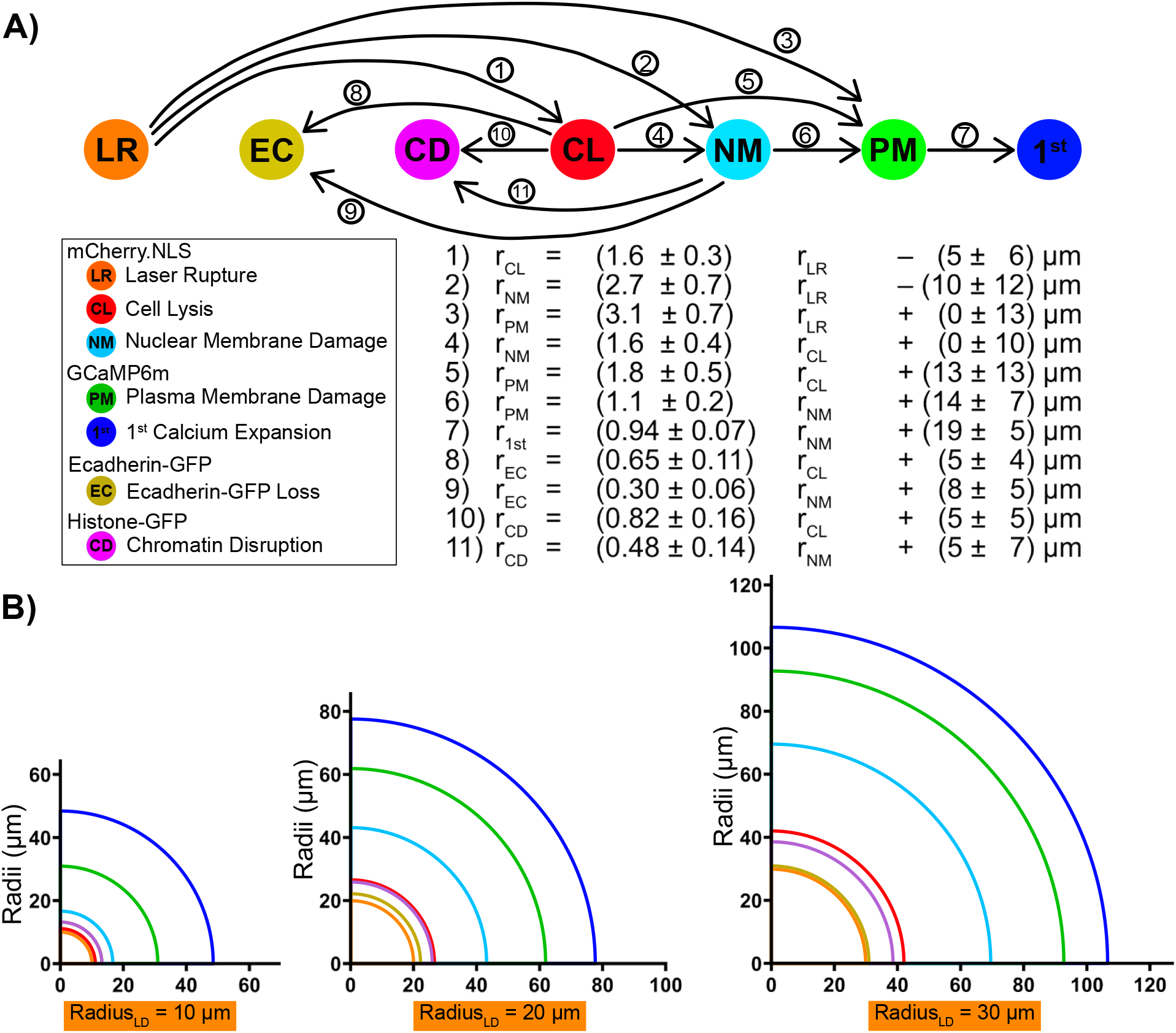
Each zone of damage can be estimated given one of them. (A) Flowchart of how to calculate each zone of damage value from a given one, with each of the eleven linear relationships displayed below corresponding to the arrows above. Arrows always go from the independent variable (x) to the dependent variable (y); the flowchart can be followed in reverse if desired. (B) Three hypothetical wounds are indicated, each with a different initial laser rupture radius (10, 20, or 30 µm). The equations in Figures 2 - 5 were used to derive the radii of the six other zones of damage from the given laser rupture radius. The 95% confidence intervals are displayed for all values. With a radius of 10 µm, the first four regions overlap (laser rupture, immediate Ecad loss, chromatin disruption, cell lysis). Because the average cell diameter in this tissue is ∼7 µm, the overlap suggests that these four regions may be within one cell diameter. As the wound gets larger, the regions become more distinguishable.

By analyzing these fluorescent proteins, we have identified many concentric zones of damage surrounding a single laser wound. These zones of damage are linearly related to one another such that we can reasonably estimate the radii of all the zones of damage, even if only one of them is known, using the equations derived in Figs. 2–5. As an example, we calculated these damage regions given a laser-rupture radius of 10, 20, or 30 µm, corresponding to small, medium, or large wounds made in this study. In doing so, we found the radii of the four inner-most regions of damage were clustered extremely tightly together in small wounds (10 and 20 µm laser rupture), such that their radii are less than a single cell diameter (∼7 µm) apart from each other. Thus, around small wounds, it is not meaningful to distinguish those four different regions of laser rupture, immediate Ecadherin loss, chromatin disruption, and cell lysis.

However, in larger wounds greater than 20 µm laser rupture, the seven regions of damage/wound signaling separate from each other and resolve into the same ascending size order: laser rupture > immediate Ecadherin loss > chromatin disruption ∼ cell lysis > nuclear membrane damage > plasma membrane damage > first calcium expansion.

## Discussion

These data show that a wound induced by pulsed laser-ablation causes tissue damage in a graded pattern and not in a uniform manner. Damage is most severe at the center of the wound, which is the focal point of the laser, and is progressively less severe with distance from the center. Importantly, cellular damage encompasses a region much larger than the area where cells are entirely destroyed, and many damaged cells recover and participate in wound repair.

From a spatial perspective, the zones of damage are arranged in a fixed order around the center. Laser wounding involves a combination of photochemical, photothermal, and photomechanical mechanism, with energy falling off with distance. A particular energy threshold likely triggers each type of damage. The first visible zone of damage is the immediate **laser rupture** around the center of the wound. At the very center of the wound, the highest laser energy causes molecular recombination where the atoms rearrange within molecules. Slightly further from the center, macromolecular assemblies are destroyed. Together these types of damage would appear as a region devoid of mCherry fluorescence in the first frame after wounding. This region of immediate laser rupture may be the same as the region of immediate **Ecad loss**; perhaps the loss of Ecadherin represents one specific type of macromolecular destruction. Although both these regions are recognized by the loss of fluorescence, we note that some molecules retain fluorescence within this area, such as histone-GFP, which retains its fluorescence even though the morphology is disrupted (discussed below).

Moving outward from the center of the wound, the next region of damage is the region of **cell lysis**, identified by the area that loses mCherry fluorescence in the minutes after wounding. We had expected cell lysis to be an immediate response to damage, but it appears instead that damage triggers lysis over the course of minutes (O’Connor et al., 2020). It is noteworthy that the region of immediate Ecadherin loss is smaller than the region of cell lysis within the first five minutes after wounding. A potential explanation is that Ecadherin is present in stable macromolecular complexes at the junctions, and therefore may appear intact even if the corresponding cell has lysed. Interestingly, Ecadherin appears to be disassembled over a much larger area during the next 90 minutes, with disassembly happening around the lysed cells and intact borders remaining only around cells that survive and participate in the wound response (Fig S1). Previous studies in embryos have reported that Ecadherin is removed from the wound margin progressively over time (Hunter et al., 2015). Our results raise the possibility that in addition to functional remodeling of the cell-cell junctions, Ecadherin removal after wounding may represent the removal of cellular debris.

The region of cell lysis largely overlaps with the region of **chromatin disruption**, assessed after lysis. It may be that chromatin appears disrupted because a cell no longer protects it; conversely, perhaps damage to the chromatin propels cell lysis. The question of whether the regions of cell lysis and chromatin disruption represent the same region brings up the issue of the resolution of detection. Although the damage profile may cause a gradient in damage to molecules, our limit of detection is a single cell, with a diameter of about 5-10 µm. Thus, two regions of damage that differ by less one-cell diameter cannot be distinguished and may have no functional difference for the tissue response to the wound. From this perspective, it may be useful to consider the regions of cell lysis and His-GFP disruption to be the same.

The next distinct zone is **nuclear membrane damage**, identified by the cloud of mCherry.NLS that leaves the nuclear compartment and floods the cytoplasm when the laser energy compromises nuclear membrane integrity. In addition to nuclei, we expect other membrane-bound organelles to be damaged by similar amounts of energy in this same region. In the more central zones of laser rupture/Ecadherin loss and the slightly larger cell lysis/His-GFP disruption, all the cells die. In contrast, in the zone of nuclear membrane damage some cells die but most cells recover and participate in wound closure (Fig. S1). Based on previous literature, it is likely that the actomyosin purse-string that surrounds the wound margin will segregate the cells that will survive and repair from those that will die.

The most distant zone of cellular damage is the zone of **plasma membrane damage**. In this region, the mechanical sheer force of the cavitation bubble rips the plasma membrane, and this zone is identified by the immediate GCaMP fluorescence from extracellular calcium flooding into the cytoplasm. In a previous study, we analyzed this zone extensively (Shannon et al., 2017) and found that it correlates with both an area of membrane depolarization and area of permeability to extracellular dyes. On kymographs, we visualized calcium entering into cells through distinct spots within this region. Cells with damaged plasma membranes in this region are able to repair the damage; indeed, the extracellular calcium is probably an important trigger for initiating plasma membrane repair (Benink and Bement, 2005; Davenport et al., 2016). As we demonstrated previously, the influx of calcium is not contained within the damaged cells, and calcium expands through gap junctions to neighboring cells in a process we call the **first calcium expansion**. Although not technically a zone of damage, this landmark is caused by damage. Interestingly, although all the other regions of damage scaled with the laser energy, the first expansion was a constant distance, about 20 µm out from the region of plasma membrane damage. This result implies that the first expansion is entirely determined by the initial influx, and the 20 µm distance is likely generated by the kinetics of the calcium buffering and re-uptake systems.

The radii of the regions of cellular damage are linearly related, in surprisingly simple relationships that we did not expect at the outset. We derived equations relating the regions of damage, and we expect these equations will be useful in estimating each region of damage when only one is known. For example, any freely diffusing fluorescent molecule expressed in cells can reveal the area of initial laser rupture and/or cell lysis, depending on the timing of data acquisition; from either of these data points, the regions of stable protein complex destruction (Ecadherin loss), nuclear membrane damage, and plasma membrane damage, and the region of high initial calcium can all be calculated.

From a temporal perspective, the initial laser pulses and following cavitation bubble cause immediate and complete cellular destruction at the very center of the wound, and simultaneously they cause a loss of nuclear membrane integrity in cells farther out, and a loss of plasma membrane integrity in cells still farther away. Next, in the milliseconds to seconds following wounding, the first known intercellular wound signal – calcium – floods into damaged cells and then travels to neighboring cells, notifying unwounded nearby cells of the damage that has just occurred. Finally, over the next minute or two, a region of cell lysis appears around the initial laser rupture, where cells lose their cytoplasmic contents.

The results of this study expand on an earlier study (Hellman et al., 2008), which identified 3 separate regions of damage after laser wounding in cell culture: a region of destruction at the center, a necrosis region of cells that die progressively over time, and a region of “molecular delivery” whereby wounded cells become permeable to the extracellular space. These seem qualitatively similar to our identified regions of laser rupture, cell lysis, and plasma membrane damage, respectively, which here we have shown in a living animal. Additionally, we report a nuclear membrane damage region and characterize sub-cellular damage markers using genetically-encoded fluorophores. Whereas Hellman, et al. analyzed how regions of damage varied with laser energy (Hellman et al., 2008), our focus is on how the regions of damage correlate with each other, with the goal of estimating the distinct regions for any pulsed-laser wound.

This study analyzed wounds made by pulsed-laser ablation, which creates reproducible and orderly wounds, with the regions of damage arranged in a reproducible manner. In trauma wounds caused by puncture, crush, or pinch wounds, all the types of damage examined in this study are expected to occur but without precise spatial patterning, rendering them difficult to identify and analyze. Thus, pulsed-laser ablation is a valuable research tool, allowing distinct regions of damage to be resolved through microscopy, and offering the opportunity to relate each type of damage with specific cellular responses.

When an epithelial barrier is breached, the cellular landscape is dramatically altered and must be repaired to avoid exposure to pathogens or loss of internal fluid (Stanisstreet, 1982; Xu and Chisholm, 2011). Although some cells at the center of a wound may be destroyed immediately or lost progressively, many partially damaged cells around the wound survive and respond by initiating the repair process, indicating that both cellular repair and tissue repair programs are activated by the same wound (Antunes et al., 2013; McNeil and Ito, 1990; Park et al., 2017). Undamaged cells farther out also receive instructive cues from the wound, recruiting them to participate in the repair process as well (Antunes et al., 2013; Danjo and Gipson, 2002; Park et al., 2017; Radice, 1980). To understand how damage initiates repair, it is critical to understand the types of damage present around a wound. However, little work has been done to characterize damaged tissue on a cellular/sub-cellular level and understand how the epithelial architecture is altered in the immediate aftermath of wounding. By characterizing zones of cellular damage, we have provided insight into what kinds of cellular changes occur to the tissue in the moments following wounding. As we have recently reported, some of these specific regions of damage initiate specific signals, and probably more have yet to be discovered. Ultimately, cellular damage itself acts as an input stimulus for the eventual behavioral output. Future studies will explore how these damage-induced cellular alterations initiate the repair process.

## Materials and Methods

### Fly lines

Drosophila crosses were maintained at 18°C for 2 days then progeny transferred to 29°C for 4-5 days before wounding.

The full genotype of each of these lines is indicated:

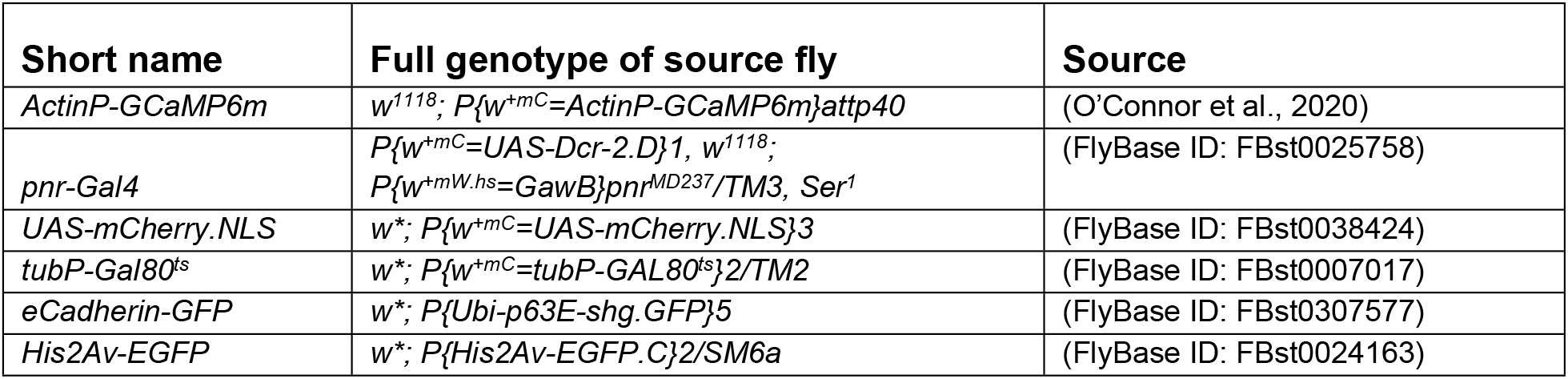

| Short name | Full genotype of source fly | Source |
| --- | --- | --- |
| *ActinP-GCaMP6m* | $w^{1118}; P\{w^{+mC}=ActinP-GCaMP6m\}attp40$ | (O'Connor et al., 2020) |
| *pnr-Gal4* | $P\{w^{+mC}=UAS-Dcr-2.D\}1, w^{1118};$<br>$P\{w^{+mW.hs}=GawB\}pnr^{MD237}/TM3, Ser^1$ | (FlyBase ID: FBst0025758) |
| *UAS-mCherry.NLS* | $w^*; P\{w^{+mC}=UAS-mCherry.NLS\}3$ | (FlyBase ID: FBst0038424) |
| *tubP-Gal80<sup>ts</sup>* | $w^*; P\{w^{+mC}=tubP-GAL80^{ts}\}2/TM2$ | (FlyBase ID: FBst0007017) |
| *eCadherin-GFP* | $w^*; P\{Ubi-p63E-shg.GFP\}5$ | (FlyBase ID: FBst0307577) |
| *His2Av-EGFP* | $w^*; P\{His2Av-EGFP.C\}2/SM6a$ | (FlyBase ID: FBst0024163) |

### Pupal mounting

Pupal mounting was performed as described previously (Shannon et al., 2017). White prepupae were identified and aged for 12-18 hours After Puparium Formation (APF) at 29°C. Multiple pupae were placed on a piece of double-sided tape (Scotch brand, catalog #665), ventral side down on a microscope slide, and their anterior pupal cases were removed with fine tipped forceps to reveal the notum epithelium (as in Fig. 1A). The entire piece of double-sided tape with dissected pupae was gently lifted from the microscope slide and adhered to a 35 mm x 50 mm coverslip (Fisherbrand, cat#125485R) so that the pupal nota were laid against the coverslip, with the pupae between the coverslip and the tape layer. Then, an oxygen permeable membrane (YSI, standard membrane kit, cat#1329882) was applied over the pupae and secured to the coverslip with additional double-sided tape so pupae would not become dehydrated or be deprived of oxygen.

### Live imaging

Live imaging of pupae was performed using a Zeiss LSM410 raster-scanning inverted confocal microscope with a 63X 1.4NA or 40X 1.3 NA oil-immersion objective. Raster-scans were performed with a 2.26 seconds scan time per image with no interval between scans.

### Laser ablation

Laser ablation was performed with the 40X objective using single pulses of the 3rd harmonic (355 nm) of a Q-switched Nd:YAG laser (5 ns pulse-width, Continuum Minilite II, Santa Clara, CA). Laser pulse energies ranged from 0.5 μJ to 10 μJ, depending on the experiment. A separate computer-controlled mirror and custom ImageJ plug-in were used to aim and operate the ablation laser so that ablation could be performed without any interruption to live imaging. The frame during ablation was retroactively considered t = 0 s.

### Calcium signal radius analysis

Calcium radius was analyzed as described previously (Shannon et al., 2017). Briefly, to quantify the spread of calcium signals from full-frame time-lapse images, the ImageJ Radial Profile Angle Plot plug-in was used on each image to determine the average GCaMP6m intensity profile as a function of distance from the center of the wound. A custom MATLAB script was then used to determine the distance from the wound at which the intensity dropped to half its maximum.

### ImageJ radius analysis

The radii of all zones of damage (apart from calcium) were found using the “measure” tool in ImageJ to find the area of a drawn circular region of interest, then deriving the radius based on A = π r^2^. The linear correlations, including confidence intervals and prediction bands, were found using Graphpad Prism.

## Acknowledgements

This work was supported by the National Institute of General Medical Sciences (1R01GM130130 to A.P.M. and M.S.H.). J.O.C. was supported by the National Institute of Child Health and Human Development (T32HD007502) and the American Heart Association (19PRE34410069 to J.O.C.). F.B.A. was supported by the Jeff Metcalf Internship Program.

## Author Contributions

Conceptualization: J.O.C., A.P.M., M.S.H; Formal Analysis: J.O.C., F.B.A.; Investigation: J.O.C.; Resources: A.P.M., M.S.H.; Data Curation: J.O.C., F.B.A.; Writing – Original Draft: J.O.C., A.P.M.; Writing – Review & Editing: J.O.C., A.P.M., M.S.H; Visualization: J.O.C., F.B.A.; Supervision: A.P.M., M.S.H; Funding Acquisition: J.O.C., F.B.A., A.P.M., M.S.H.

## Supplementary Data

**Figure S1.**
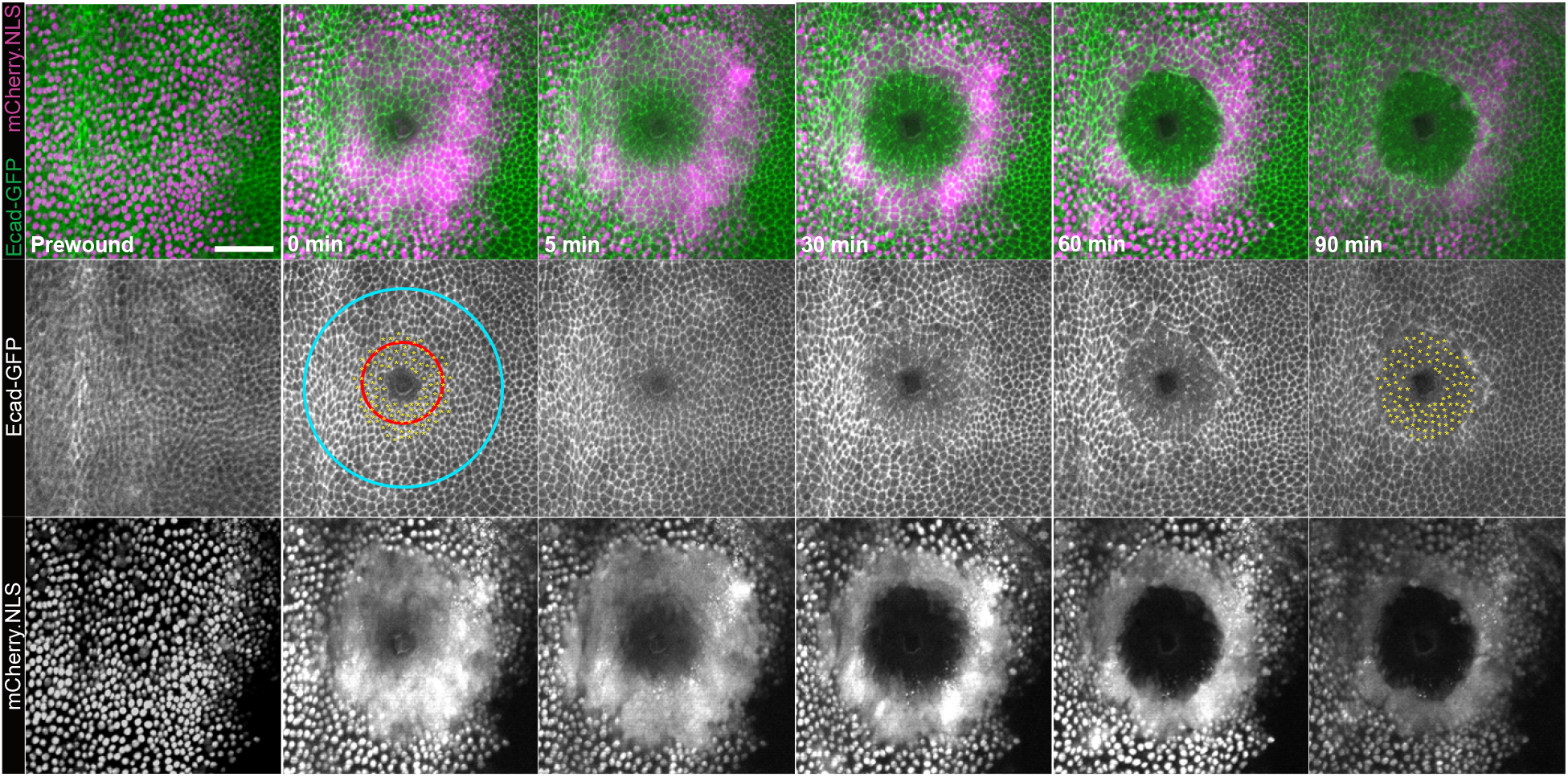
Timelapse of Ecadherin-GFP (green) and pnr>mCherry.NLS (magenta) following wounding, from Movie 1. Immediately after wounding (0 min), Ecad is lost from the center of the wound, as analyzed in Fig. 4; by 90 minutes, the region of of Ecad loss is much larger. During these ∼90 minutes, Ecad appears to be gradually lost evenly across the region of cell lysis (red circle) where mCherry.NLS is lost. Cells that have Ecad borders at 0 min but eventually lose Ecad (labeled with yellow asterisks) clearly die in response to wounding. The region of nuclear membrane damage (blue circle) is evident by mCherry.NLS escaping from the nucleus immediately after wounding. Some cells with nuclear membrane damage to recover whereas those closer to the center die. Scale bar = 50 µm.

### Movie 1

Timelapse movie of Ecadherin-GFP (green) and pnr>mCherry.NLS (magenta) over 90 minutes following wounding. Movie shows Ecadherin gradual loss in cells that initially appeared intact.

## References

Antunes, M., Pereira, T., Cordeiro, J.V., Almeida, L., and Jacinto, A. (2013). Coordinated waves of actomyosin flow and apical cell constriction immediately after wounding. The Journal of Cell Biology 202, 365–379.

Benink, H.A., and Bement, W.M. (2005). Concentric zones of active RhoA and Cdc42 around single cell wounds. The Journal of Cell Biology 168, 429–439.

Bosch, M., Serras, F., Martín-Blanco, E., and Baguñà, J. (2005). JNK signaling pathway required for wound healing in regenerating Drosophila wing imaginal discs. Developmental Biology 280, 73–86.

Brand, A.H., and Perrimon, N. (1993). Targeted gene expression as a means of altering cell fates and generating dominant phenotypes. Dev Camb Engl 118, 401–415.

Burra, S., Wang, Y., Brock, A.R., and Galko, M.J. (2013). Wound Regeneration and Repair, Methods and Protocols. Methods Mol Biology 1037, 449–461.

Capilla, A., Karachentsev, D., Patterson, R.A., Hermann, A., Juarez, M.T., and McGinnis, W. (2017). Toll pathway is required for wound-induced expression of barrier repair genes in the Drosophila epidermis. Proc National Acad Sci 114, E2682–E2688.

Carvalho, L., Jacinto, A., and Matova, N. (2014). The Toll/NF-κB signaling pathway is required for epidermal wound repair in Drosophila. Proceedings of the National Academy of Sciences of the United States of America 111, E5373–82.

Danjo, Y., and Gipson, I.K. (2002). Specific Transduction of the Leading Edge Cells of Migrating Epithelia Demonstrates That They are Replaced During Healing. Exp Eye Res 74, 199–204.

Davenport, N.R., Sonnemann, K.J., Eliceiri, K.W., and Bement, W.M. (2016). Membrane dynamics during cellular wound repair. Molecular Biology of the Cell 27, 2272–2285.

Enyedi, B., and Niethammer, P. (2015). Mechanisms of epithelial wound detection. Trends Cell Biol 25, 398–407.

Fauvarque, M.-O., and Williams, M.J. (2011). Drosophila cellular immunity: a story of migration and adhesion. J Cell Sci 124, 1373–1382.

Galko, M.J., and Krasnow, M.A. (2004). Cellular and Genetic Analysis of Wound Healing in Drosophila Larvae. Plos Biol 2, e239.

Gault, W.J., Enyedi, B., and Niethammer, P. (2014). Osmotic surveillance mediates rapid wound closure through nucleotide release. J Cell Biol 207, 767–782.

Hellman, A.N., Rau, K.R., Yoon, H.H., and Venugopalan, V. (2008). Biophysical Response to Pulsed Laser Microbeam-Induced Cell Lysis and Molecular Delivery. J Biophotonics 1, 24–35.

Hunter, M.V., Lee, D.M., Harris, T.J.C., and Fernandez-Gonzalez, R. (2015). Polarized E-cadherin endocytosis directs actomyosin remodeling during embryonic wound repair. The Journal of Cell Biology 210, 801–816.

Hunter, M.V., Willoughby, P.M., Bruce, A.E.E., and Fernandez-Gonzalez, R. (2018). Oxidative Stress Orchestrates Cell Polarity to Promote Embryonic Wound Healing. Developmental Cell 47, 377-387.e4.

Hutson, M.S., and Ma, X. (2007). Plasma and Cavitation Dynamics during Pulsed Laser Microsurgery in vivo. Phys Rev Lett 99, 158104.

Leeuwen, T.G.V., Jansen, E.D., Motamedi, M., Borst, C., and Welch, A.J. (1995). Optical-Thermal Response of Laser-Irradiated Tissue. 709–763.

Luster, A.D., Alon, R., and Andrian, U.H. von (2005). Immune cell migration in inflammation: present and future therapeutic targets. Nat Immunol 6, 1182–1190.

Martin, P., and Lewis, J. (1992). Actin cables and epidermal movement in embryonic wound healing. Nature 360, 179–183.

Martin, P., Nobes, C., McCluskey, J., and Lewis, J. (1994). Repair of excisional wounds in the embryo. Eye 8, 155–160.

Matsubayashi, Y., and Millard, T.H. (2016). Analysis of the Molecular Mechanisms of Reepithelialization in Drosophila Embryos. Adv Wound Care 5, 243–250.

McNeil, P.L., and Ito, S. (1989). Gastrointestinal Cell Plasma Membrane Wounding and Resealing In Vivo. Gastroenterology 96, 1238–1248.

McNeil, P.L., and Ito, S. (1990). Molecular traffic through plasma membrane disruptions of cells in vivo. J Cell Sci 96 (Pt 3), 549–556.

McNeil, P.L., and Steinhardt, R.A. (1997). Loss, Restoration, and Maintenance of Plasma Membrane Integrity. J Cell Biology 137, 1–4.

O’Connor, J.T., Stevens, A.C., Shannon, E.K., Akbar, F.B., LaFever, K.S., Narayanan, N., Hutson, M.S., and Page-McCaw, A. (2020). A protease-initiated model of wound detection. Biorxiv 2020.12.08.415554.

Park, S., Gonzalez, D.G., Guirao, B., Boucher, J.D., Cockburn, K., Marsh, E.D., Mesa, K.R., Brown, S., Rompolas, P., Haberman, A.M., et al. (2017). Erratum: Tissue-scale coordination of cellular behaviour promotes epidermal wound repair in live mice. Nat Cell Biol 19, 407–407.

Radice, G.P. (1980). The spreading of epithelial cells during wound closure in Xenopus larvae. Dev Biol 76, 26–46.

Rämet, M., Lanot, R., Zachary, D., and Manfruelli, P. (2002). JNK signaling pathway is required for efficient wound healing in Drosophila. Developmental Biology 241, 145–156.

Ramos-Lewis, W., LaFever, K.S., and Page-McCaw, A. (2018). A scar-like lesion is apparent in basement membrane after wound repair in vivo. Matrix Biol 74, 101–120.

Razzell, W., Evans, I.R., Martin, P., and Wood, W. (2013). Calcium flashes orchestrate the wound inflammatory response through DUOX activation and hydrogen peroxide release. Current Biology : CB 23, 424–429.

Shabir, S., and Southgate, J. (2008). Calcium signalling in wound-responsive normal human urothelial cell monolayers. Cell Calcium 44, 453–464.

Shannon, E.K., Stevens, A., Edrington, W., Zhao, Y., Jayasinghe, A.K., Page-McCaw, A., and Hutson, M.S. (2017). Multiple Mechanisms Drive Calcium Signal Dynamics around Laser-Induced Epithelial Wounds. Biophysical Journal 113, 1623–1635.

Stanisstreet, M. (1982). Calcium and wound healing in Xenopus early embryos. Journal of Embryology and Experimental Morphology 67, 195–205.

Stramer, B., Wood, W., Galko, M.J., Redd, M.J., Jacinto, A., Parkhurst, S.M., and Martin, P. (2005). Live imaging of wound inflammation in Drosophila embryos reveals key roles for small GTPases during in vivo cell migration. The Journal of Cell Biology 168, 567–573.

Thuma, L., Carter, D., Weavers, H., and Martin, P. (2018). Drosophila immune cells extravasate from vessels to wounds using Tre1 GPCR and Rho signaling. The Journal of Cell Biology 217, 3045–3056.

Troutman, R.C., Véronneau-Troutman, S., Jakobiec, F.A., and Krebs, W. (1986). A new laser for collagen wounding in corneal and strabismus surgery: a preliminary report. T Am Ophthal Soc 84, 117–132.

Venugopalan, V., Guerra, A., Nahen, K., and Vogel, A. (2002). Role of Laser-Induced Plasma Formation in Pulsed Cellular Microsurgery and Micromanipulation. Phys Rev Lett 88, 078103.

Welch, A.J., Motamedi, M., Rastegar, S., LeCarpentier, G.L., and Jansen, D. (1991). Laser Thermal Ablation. Photochem Photobiol 53, 815–823.

Wood, W., Jacinto, A., Grose, R., Woolner, S., Gale, J., Wilson, C., and Martin, P. (2002). Wound healing recapitulates morphogenesis in Drosophila embryos. Nature Cell Biology 4, 907–912.

Wu, Y., Brock, A.R., Wang, Y., Fujitani, K., Ueda, R., and Galko, M.J. (2009). A blood-borne PDGF/VEGF-like ligand initiates wound-induced epidermal cell migration in Drosophila larvae. Current Biology : CB 19, 1473–1477.

Xu, S., and Chisholm, A.D. (2011). A Gαq-Ca2+ signaling pathway promotes actin-mediated epidermal wound closure in C. elegans. Current Biology : CB 21, 1960–1967.

Yoo, S.K., Freisinger, C.M., LeBert, D.C., and Huttenlocher, A. (2012). Early redox, Src family kinase, and calcium signaling integrate wound responses and tissue regeneration in zebrafish. The Journal of Cell Biology 199, 225–234.

